# [^225^Ac]Ac/[^89^Zr]Zr-labeled N4MU01 radioimmunoconjugates as theranostics against nectin-4 positive triple negative breast cancer

**DOI:** 10.1101/2024.03.04.583420

**Authors:** Hanan Babeker, Fabrice Ngoh Njotu, Jessica Pougoue Ketchemen, Anjong Florence Tikum, Alireza Doroudi, Emmanuel Nwangele, Maruti Uppalapati, Humphrey Fonge

## Abstract

**Purpose:** Nectin-4 is an overexpressed biomarker in 60-70% of triple-negative breast cancer (TNBC), and an ideal target for radiotherapy and PET imaging. In this study, we have developed theranostic radioimmunoconjugates (RICs) based on a fully-human anti-nectin-4 antibody, N4MU01. We characterized and evaluated the efficacy of these RICs for applications in molecular imaging and radiotherapy of aggressive TNBC models.

**Methods:** An anti-nectin-4 antibody (N4MU01) was radiolabeled with ^89^Zr and ^225^Ac, for imaging and radiotherapy, respectively, using TNBC xenograft and syngeneic models. Biodistribution & PET imaging of [^89^Zr]Zr-DFO-N4MU01 RIC was studied in mice bearing nectin-4 positive xenografts. Dosimetry of [^225^Ac]Ac-Macropa-N4MU01 was studied in healthy mice, and therapeutic efficacy was evaluated in mice bearing human TNBC MDA-MB-468 xenograft and in a syngeneic xenograft model using murine 4T1 breast cancer cells transfected with human nectin-4 (4T1._nectin-4_). Mice received 2 doses of 13 kBq or 18.6 kBq 10 days apart.

**Results:** The pharmacokinetic profile of [^89^Zr]Zr-DFO-N4MU01 RIC showed biphasic distribution with a moderate elimination half-life of 63 h. PET imaging and biodistribution of [^89^Zr]Zr-DFO-N4MU01 in mice bearing MDA-MB-468 xenograft showed high tumor uptake of 13.2 ± 1.12 %IA/g at 120 h. [^225^Ac]Ac-Macropa-N4MU01 was effectively internalized in MDA-MB-468 and was cytotoxic to the cells with IC_50_ of 1.2 kBq/mL. Mice bearing MDA-MB-468 xenograft treated with [^225^Ac]Ac-Macropa-N4MU01 (13 kBq or 18.6 kBq) had a remarkable tumor growth inhibition that was dose dependent. For the syngeneic 4T1._nectin-4_ model, treatment with [^225^Ac]Ac-Macropa-N4MU01 (13 kBq) led to complete tumor remission in 83.3% (5/6) of mice.

**Conclusion:** The specific tumor uptake and remarkable effectiveness at shrinking aggressive TNBC tumors is very promising towards clinical development of N4MU01 RICs as theranostics against TNBC.

## Introduction

Breast cancer (BC) is the most common cancer in women, with 1 in 8 women likely to be diagnosed with the disease in their lifetime. Triple-negative breast cancer (TNBC) represents 10%– 20% of invasive BC and has been associated with African-American race, deprivation status, younger age at diagnosis, more advanced disease stage at diagnosis, higher grade, high mitotic indices, family history of breast cancer and breast cancer gene 1 (BRCA1) mutations, and is aggressive with poor prognosis [1]. Taxanes and anthracyclines alone, or in combination constitute the first line of treatment for metastatic BC with response rates of 38% and 46% for single agent and combination-based therapies, respectively [2]. However, *de novo* resistance to taxanes is common in over 50% of patients and most patients acquire resistance [2]. Sacituzumab govitecan (SG) is the only antibody-based therapeutics recently approved for the treatment of TNBC. SG is an anti-TROP2 antibody-drug conjugate (ADC) in which the anti-TROP2 antibody is conjugated to cytotoxic drug SN38, and it shows significantly improved benefits compared with chemotherapy [3]. The objective response rate (ORR) was just 33% for SG (complete response 4%, partial response (PR) 27%) compared with 4% (all PR) for chemotherapy [3]. Despite this promising outcome, novel approaches are required.

The poliovirus receptor-related protein 4 (PVRL4/nectin-4) is uniquely overexpressed in most cancers of epithelial origin (including gastric, breast, lung, pancreatic ovarian, bladder and esophageal) but not in non-malignant or healthy tissues [4, 5]. Since its identification less than two decades ago, nectin-4 has become a highly sought-after target in precision oncology. Nectin-4 is a member of the nectin family of immunoglobulin-like adhesion molecules that are proposed to mediate Ca^2+^-independent cell–cell adhesion via both homophilic and heterophilic trans-interactions at adherens junctions where they recruit cadherins and modulate cytoskeleton rearrangements [6]. Notably, large clinical studies have consistently confirmed 60 – 70% of all TNBC overexpress nectin-4 [5, 7, 8]. Immunohistochemical analysis of a panel of 294 healthy tissue specimens from 36 human (healthy) organs showed very weak homogenous staining mainly in human skin keratinocytes, skin appendages (sweat glands and hair follicles), transitional epithelium of bladder, salivary gland (ducts), esophagus, breast, and stomach, with no staining from other tissues [8]. Nectin-4 plays an important role in tumor cell proliferation, angiogenesis, migration, and metastasis through PI3K/Akt signaling pathway [9–14]. Therefore, nectin-4 is an excellent biomarker for targeted radioimmunotherapy of TNBC.

Enfortumab vedotin (PADCEV^®^) is a fully human anti-nectin-4 antibody drug conjugate in which the antibody is conjugated to monomethyl auristatin (MMAE) via a cleavable linker, and is approved for the treatment of nectin-4 positive advanced urothelial cancers [15]. In preclinical studies enfortumab vedotin was effective in mice bearing nectin-4 positive TNBC tumors, and is currently in phase II trials in patients with advanced metastatic solid tumors including TNBC (EV-202 trial) [7]. In mice bearing nectin-4 positive TNBC xenografts, complete remission was seen in most tumors but unfortunately all re-grew after periods of no-treatment. Although these recurrent tumors were sensitive to the drug, the same manner of tumor re-growth was observed after subsequent secession of treatment [7]. Recently, Cabaud *et al* [16] showed preclinically that the mechanism of resistance to anti-nectin-4 ADC was due to the expression of ABCB1gene, encoding the multidrug resistance protein MDR1/P-glycoprotein (P-gp), associated with focal gene amplification and high protein expression. This appears to be a common mechanism of resistance of most ADCs and this mirrors that of chemotherapeutics [16–18]. Unlike ADCs, resistance to alpha particle therapeutics has never been observed [19, 20]. Alpha particle therapeutics are up to 1000-fold more potent than beta-emitting therapeutics, are effective in both normoxic and hypoxic environments. The characteristics of ^225^Ac: t_1/2_ 10.0 days; energy range of 4 αs is 6–8 MeV (cumulatively 28 MeV/decay) decays with the emission of 4 αs with a range of 50–80 *μ*m and linear energy transfer (LET) of 0.16 MeV/μm make it very potent for the treatment of TNBC. We have previously developed a fully human anti-nectin-4 (N4MU01) antibody using phage display (parallel manuscript). In this work, we developed anti-nectin-4 theranostics (therapeutics and diagnostics) based on N4MU01. N4MU01 diagnostic agent was developed by conjugating the antibody with deferoxamine (using *p*-SCN-Bn-DFO) and radiolabeling with zirconium-89 (^89^Zr) to obtain a PET imaging agent. N4MU01 radiotherapeutic was developed by conjugating the antibody with 18-membered ring macrocyclic chelator Macropa (using *p*-SCN-Macropa) followed by radiolabeling with actinium-225 (^225^Ac) for alpha particle therapy.

## Materials and methods

### Cell lines and xenografts

Human breast cancer cell lines MDA-MB-468, MCF-7 which expresse nectin-4, and MDA-MB-231, known for nectin-4 negative expression, were purchased from ATCC (Rockville, MD). For syngeneic mouse model, mouse breast cancer cell line 4T1 was purchased from ATCC (Rockville, MD) and transfected with full-length nectin-4 protein (4T1._nectin-4_). Cell culture conditions and xenograft establishment is provided in the supplementary material. Animals used in this study were maintained following the guidelines of the University of Saskatchewan Animal Care Committee (protocol # 20170084 and 20220021). Cell lines were authenticated using short tandem repeat (STR) profiling (Centre for Applied Genomics, Hospital for SickKids, Toronto, ON) and had no detectable mycoplasma prior to their use.

### 4T1 cell transfection with human nectin-4

The human nectin-4 coding sequence was amplified by PCR, and the PCR product was cloned into a custom lentiviral vector pMUFV01 linearized after digestion with enzymes (EcoRI and NheI) using Gibson assembly. The fully assembled vector map is shown in Supplementary Figure 1. The complete method of transfection and sorting of the cells is in the supplementary methods section.

### Characterization of N4MU01 binding using flow cytometry

The binding affinity and specificity of N4MU01 were evaluated by flow cytometry using nectin-4 expressing breast cancer cells MDA-MB-468, MCF-7, 4T1._nectin-4_ and nectin-4 negative MDA-MB-231 cells as described previously [21]. The detailed method is described in the supplementary section.

### Internalization of N4MU01

The internalization of N4MU01 was studied for 48 h using the Incucyte live-cell imaging system as described previously [21]. The detailed method is described in the supplementary method section.

### Conjugation with bifunctional chelators and radiolabeling with ^89^Zr and ^225^Ac

N4MU01 was conjugated with *p*-SCN-Bn-deferoxamine (DFO) for labeling with ^89^Zr as described previously [22]. Quality control of the immunoconjugate was done using size exclusion chromatography (SEC) high-performance liquid chromatography (HPLC) (SEC-HPLC), flow cytometry, and automated electrophoresis (2100 Bioanalyzer, Agilent, Santa Clara, CA, USA). Radiolabeling of DFO-N4MU01 with ^89^Zr and purification were done as reported previously [22].

18-membered macrocyclic bifunctional chelator 6-((16-((6-carboxypyridin-2-yl)methyl)-1,4,10,13-tetraoxa-7,16-diazacyclooctadecan-7-yl)methyl)-4-isothiocyanatopicolinic acid (*p*-SCN-Macropa) was synthesized as reported earlier [23]. The conjugation of N4MU01 with Macropa for the labeling with ^225^Ac was performed following the lab SOP (supplementary methods). Quality control of the immunoconjugate was done using SEC-HPLC, matrix-assisted laser desorption (MALDI) time of flight (TOF) (MALDI-TOF) and flow cytometry. Radiolabeling with ^225^Ac was performed as previously reported [22].

The stability of [^89^Zr]Zr-DFO-labeled and [^225^Ac]Ac-Macropa-labeled N4MU01 conjugates at 37 °C was investigated using instant thin-layer chromatography (iTLC) by analyzing aliquots of the radiolabeled over time. [^89^Zr]Zr-DFO-labeled or [^225^Ac]Ac-Macropa-labeled conjugates were mixed with human plasma or 1x PBS solution at a final concentration of 15 MBq/mL and 2.5 MBq/mL, respectively and incubated at 37 °C. Aliquots of the solution were taken every 24 h for five days and analyzed for radiochemical purity using iTLC.

### Radioligand binding assay of [^89^Zr]Zr-DFO-N4MU01 and [^225^Ac]**Ac**-Macropa–N4MU01

The binding of [^89^Zr]Zr-DFO-N4MU01 to nectin-4 positive MCF-7 cells and [^225^Ac]Ac-Macropa-N4MU01 to nectin-4 positive MDA-MB-468 was determined using a saturation radioligand binding assay as described previously [24]. The detailed method is described in the supplementary method section.

### MicroPET/CT imaging and biodistribution of [^89^Zr]Zr-DFO-N4MU01

Female CD-1 nude mice (n = 4) bearing nectin-4 positive MDA-MB-468 xenograft and female BALB/c mice (n=4) bearing 4T1._nectin-4_ tumors for syngeneic mouse model were injected intravenously using 12 ± 1 MBq (21 – 26 µg) [^89^Zr]Zr-DFO-N4MU01. Additionally, mice bearing MDA-MB-468 xenograft (n = 4) were injected intravenously with 200 µg of unlabeled N4MU01 four hours prior to the same injection of [^89^Zr]Zr-DFO-N4MU01 to pre-block nectin-4 receptors. MicroPET/CT imaging was done using the Vector^4^CT scanner at different times. Biodistribution studies involved sacrificing mice at 24 and 120 h post-injection (p.i.), with radioactivity measured in all major organs and blood. The detailed methods are in the supplementary materials.

### Pharmacokinetic profile of [^89^Zr]Zr-DFO-N4MU01

The pharmacokinetics of [^89^Zr]Zr-DFO-N4MU01 were studied in healthy female CD-1 nude mice (n = 3/group) as described previously [25]. The detailed methods are in the supplementary materials.

### *In vitro* cytotoxicity

The *in vitro* cytotoxicity (IC_50_ values) of [^225^Ac]Ac-Macropa-N4MU01, control immunoconjugates anti-CD20 [^225^Ac] Ac-Macropa-rituximab, and unlabeled N4MU01 in nectin-4 positive MDA-MB-468, MCF-7, and nectin-4 negative MDA-MB-231 was determined using IncuCyte Cytotox Red reagent in an IncuCyte S3 live-cell imager (Essen BioScience, AnnArbor, MI) as previously reported [24]. The detailed methods are in the supplementary materials.

### Biodistribution and dosimetry of [^225^Ac]Ac-Macropa-N4MU01

To estimate radiation dose to healthy tissues, normal BALB/c nude mice (n ≥ 4/group) were administered 13 kBq of [^225^Ac]Ac-Macropa-N4MU01 via tail vein and sacrificed at 1, 24, 48, 120, and 264 h p.i. followed by biodistribution studies. The mouse biodistribution % injected activity per gram (%IA/g) data was extrapolated to human data (%IA) using the formula % IA (human) = % IA/g (mouse) x total body weight of mouse (in kg) x mass of human organ (in g) / total body weight of human (in kg). For each organ, this was plotted against sampling time and used to obtain an estimate of the residence time of the agent in the organ in MBq-h/MBq, represented by the area under the time-activity function integrated to infinity (complete decay) of the ^225^Ac. The residence time was fitted into the OLINDA kinetics model (OLINDA/EXM V2.2, Hermes Medical Solutions Montreal QC) to generate absorbed doses in units of cGy (centi-gray) per millicurie (cGy/mCi) of ^225^Ac administered [29, 30].

### [^225^Ac]Ac-Macropa-N4MU01 radioimmunotherapy

*In vivo* efficacy studies were done using athymic nude mice and immune-competent BALB/c mice aged 4 – 6 weeks bearing MDA-MB-468 and 4T1._nectin-4_ tumors, respectively. All the experiments and euthanasia were performed in accordance with UACC guidelines. Mice bearing MDA-MB-468 xenograft were divided into 5 groups (n ≥ 4/group), namely, [^225^Ac]Ac-Macropa-N4MU01 (two doses of 13 kBq /dose), [^225^Ac]Ac-Macropa-N4MU01 (two doses of 18.6 kBq /dose), saline treatment, and unlabeled N4MU01. Whereas mice bearing 4T1._nectin-4_ tumors were divided into 4 groups (n ≥ 5/group), namely, [^225^Ac]Ac-Macropa-N4MU01 (two doses of 13 kBq /dose), therapeutic dose of N4MU01 (two doses of 100 μg /dose), N4MU01 (two doses of 1.3 μg/dose), and saline control. All treated mice received two treatment doses via a tail vein on days 0 and 10. Tumor growth was monitored by measuring the greatest length and width using a digital caliper (tumor volume = (length × width^2^)/2). The study was terminated when tumors reached a volume ≥ 1500 mm^3^. These volumes were used to determine survival in the different groups using Kaplan-Meier curves. The body weight of each mouse was recorded during the experimental period.

### Statistical analysis

All data were expressed as the mean ± standard deviation or mean ± standard error of mean of at least three independent experiments. A two-tailed Student’s *t-*test or analysis of variance (ANOVA) with Bonferroni post hoc test was used to assess the statistical significance between the groups. All graphs were prepared and analyzed using GraphPad Prism (version 9; GraphPad, La Jolla, CA, USA) and a *p*-value ≤ of 0.05 was considered significant.

## Results

### *In vitro* characterization of N4MU01

*In vitro* characterization using flow cytometry was performed to determine the binding of N4MU01 to nectin-4 positive MDA-MB-468, MCF-7, 4T1 cells transfected with human nectin-4 (4T1._nectin-4_) (Supplementary Figure 1), and nectin-4 negative MDA-MB-231 cell lines. Dose-dependent specific binding was observed with MDA-MB-468, MCF-7, 4T1._nectin-4_ and not MDA-MB-231 cells (Figure 1A).

**Figure 1:**
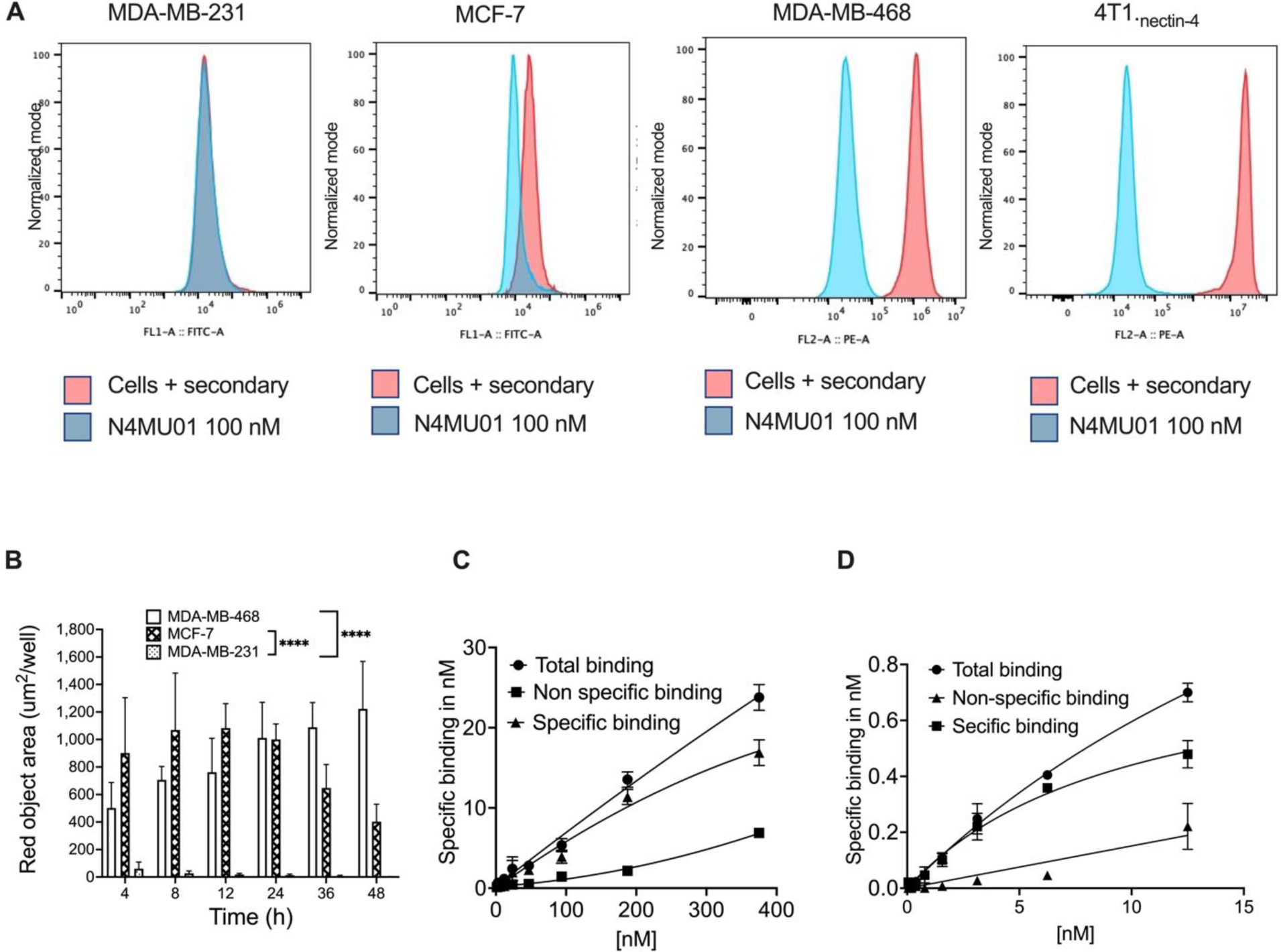
Flow cytometry binding, internalization and radioligand binding assay of N4MU01 and its conjugates. Different concentrations of N4MU01 were incubated with **A**) MDA-MB-231, MCF-7, MDA-MB-468, and 4T1._nectin-4_ cells. Dose-dependent binding was observed in nectin-4 expressing cell lines, MDA-MB-468, MCF7, and 4T1._nectin-4_. No binding was observed in MDA-MB-231 control cell line. **B**) Internalization of N4MU01: MDA-MB-468, MCF-7, and MDA-MB-231 cells were treated with IncuCyte FabFluor-labeled N4MU01 (4 μg/mL); HD phase and red fluorescence images (10×) were captured every 2 h for 48 h. All data are shown as a mean ± SEM (n=3), and the average internalization values for MCF-7 and MDA-MB-468 were compared with that of MDA-MB-231 using ANOVA. C) Estimation of K_D_ values using radioligand binding assay for C ) [^89^Zr]Zr-DFO-N4MU01 in nectin-4 positive MCF-7 and D) [^225^Ac]-Macropa-N4MU01 in nectin-4 positive MDA-MB-468 cells. \*\*\*\**p* < 0.0001.

### N4MU01 internalization

A time-dependent increase in red fluorescence was observed in nectin-4 expressing MDA-MB-468 (high expression) and MCF-7 (moderate expression) cell lines, respectively, but not in the negative control MDA-MB-231 cells at 4 – 48 h post incubation (Figure 1B). The N4MU01 showed 72- and 1100-fold higher internalization in nectin-4 positive MDA-MB-468 cells compared with the negative control MDA-MB-231 cells at 24 and 48 h, respectively. In MCF-7 cells, the N4MU01 showed 71 and 365-fold higher internalization compared to the negative control MDA-MB-231 cells.

### Conjugation and quality control of N4MU01 radioimmunoconjugates

HPLC analysis showed that the conjugation of *p*-SCN-DFO to N4MU01 resulted in > 99% pure DFO-N4MU01 with no aggregates. The size and purity of DFO-N4MU01 were evaluated using a bioanalyzer (Supplementary Figure 2A & 2B). Similarly, the conjugation of Macropa to N4MU01 resulted in >99% pure immunoconjugate with no aggregates (Supplementary Figure 2C & 2D). Automated electrophoresis using bioanalyzer showed that DFO-N4MU01 was 90% pure with a molecular weight of 153.2 kDa (vs 150.2 kDa for unconjugated N4MU01). This indicates that there were 4.0 DFO chelator molecules/antibody molecule.

*In vitro* saturation binding assay of unconjugated N4MU01 and DFO-N4MU01 was done using nectin-4 positive MCF-7 cells using flow cytometry. The estimated K_D_ values for unconjugated N4MU01 and DFO-N4MU01 were 2.9 and 3 nM, respectively. The estimated EC_50_ values for unconjugated N4MU01 and DFO-N4MU01 were 7.3 and 20.0 nM, respectively (Supplementary Figure 3 ).

N4MU01 conjugated with DFO or Macropa were labeled with ^89^Zr or ^225^Ac, respectively. The radiochemical yield (RCY) of [^89^Zr]Zr-DFO-N4MU01 was > 90% at a specific activity of 0.5 MBq/μg. Similarly, the RCY of [^225^Ac]Ac-Macropa-N4MU01 was > 95% at a specific activity of 10 kBq/μg (Supplementary Figure 2C & 2D). The stability of [^89^Zr]Zr-DFO-N4MU01 and [^225^Ac]-Ac-Macropa-N4MU01 was determined at different time points at 37 °C in PBS and human plasma using iTLC. More than 95% of [^89^Zr]Zr-DFO-N4MU01 remained intact for 96 h in human serum and for 72 h in PBS. Similarly, more than 96% of [^225^Ac]Ac-Macropa-N4MU01 remained intact for 120 h in human serum and was ≥ 91% stable in PBS (Supplementary Figure 4).

Saturation radioligand binding assay using [^89^Zr]Zr-DFO-N4MU01 on MCF7 cells and [^225^Ac]Ac-Macropa-N4MU01 on MDA-MB-468 showed dose-dependent increase in specific binding (Figure 1C and 1D). The estimated K_D_ of [^89^Zr]Zr-DFO-N4MU01 was 10 nM which is 3.4-fold lower affinity than unlabeled N4MU01 and 3.3-fold lower affinity than the DFO-N4MU01. The estimated K_D_ of [^225^Ac]Ac-Macropa-N4MU01 was 14.8 nM which is 3.7-fold lower affinity than unlabeled N4MU01 and similar to the Macropa-N4MU01.

### Pharmacokinetics, biodistribution, and microPET/CT of [^89^Zr]Zr-DFO-N4MU01

Pharmacokinetic profile of [^89^Zr]Zr-DFO-N4MU01 injected in CD-1 nude mice showed fast distribution half-life t_1/2α_ of 3.4 h and a moderate elimination t_1/2ß_ of 63 h (Figure 2).

**Figure 2:**
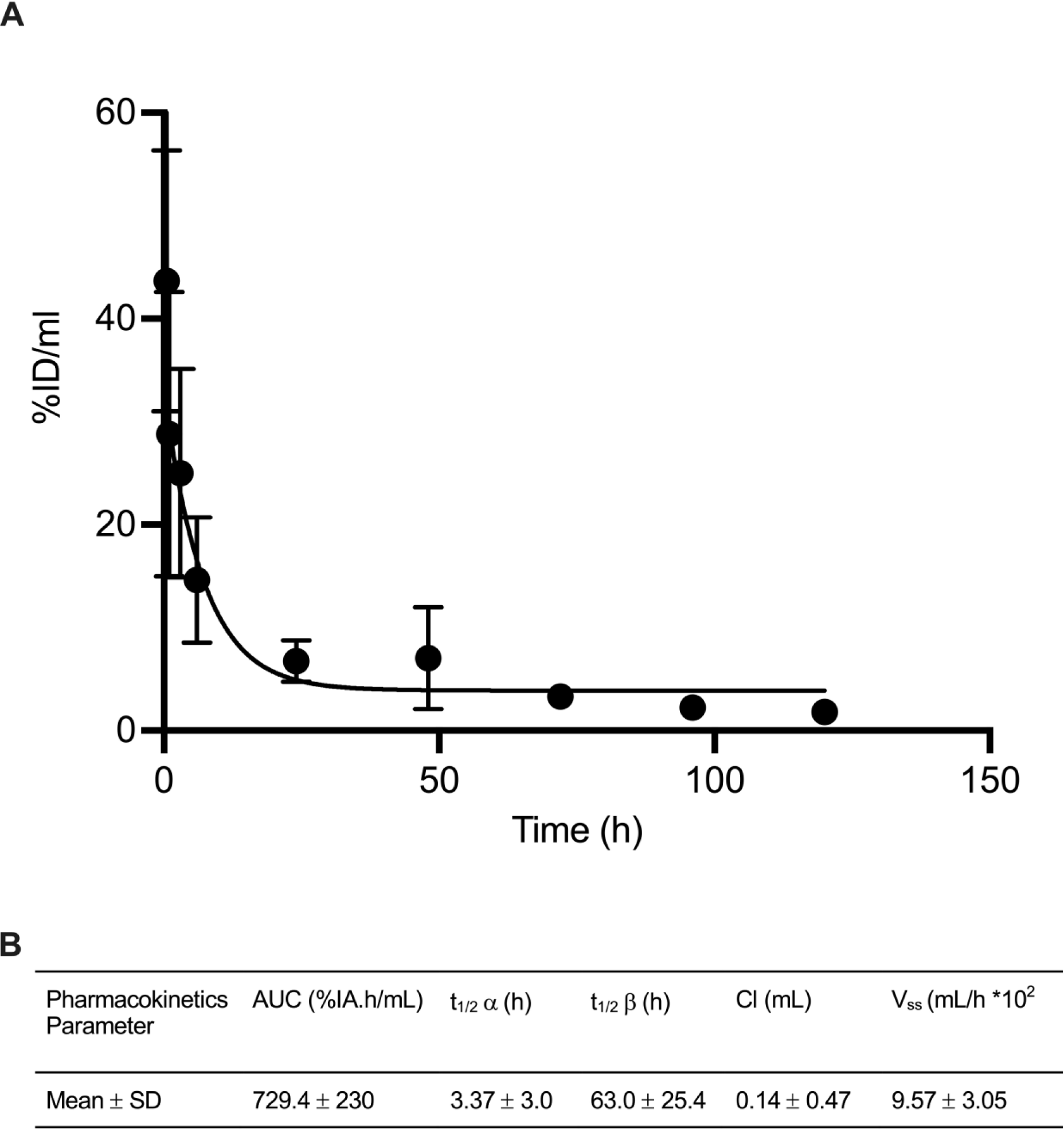
Pharmacokinetics of [^89^Zr]Zr-DFO-N4MU01 in healthy CD-1 nude mice. **A**) [^89^Zr]Zr-DFO-N4MU01 showed a bi-phasic half-life with **B**) a fast distribution and moderate elimination.

MicroPET/CT imaging experiments were conducted with [^89^Zr]Zr-DFO-N4MU01 in mice bearing MDA-MB-468 xenograft and in syngeneic mouse model bearing 4T1_nectin-4_ tumor. High tumor uptake was delineated in nectin-4 positive xenografts as shown by microPET/CT imaging at 24 – 120 h p.i. (Figure 3A). There was visibly very low tumor uptake of [^89^Zr]Zr-DFO-N4MU01 when pre-blocked with unlabeled antibody (Figure 3A). For the syngeneic mouse model, tumor uptake increased over time, as shown by microPET/CT imaging at 24 and 120 h p.i (Figure 3B).

**Figure 3:**
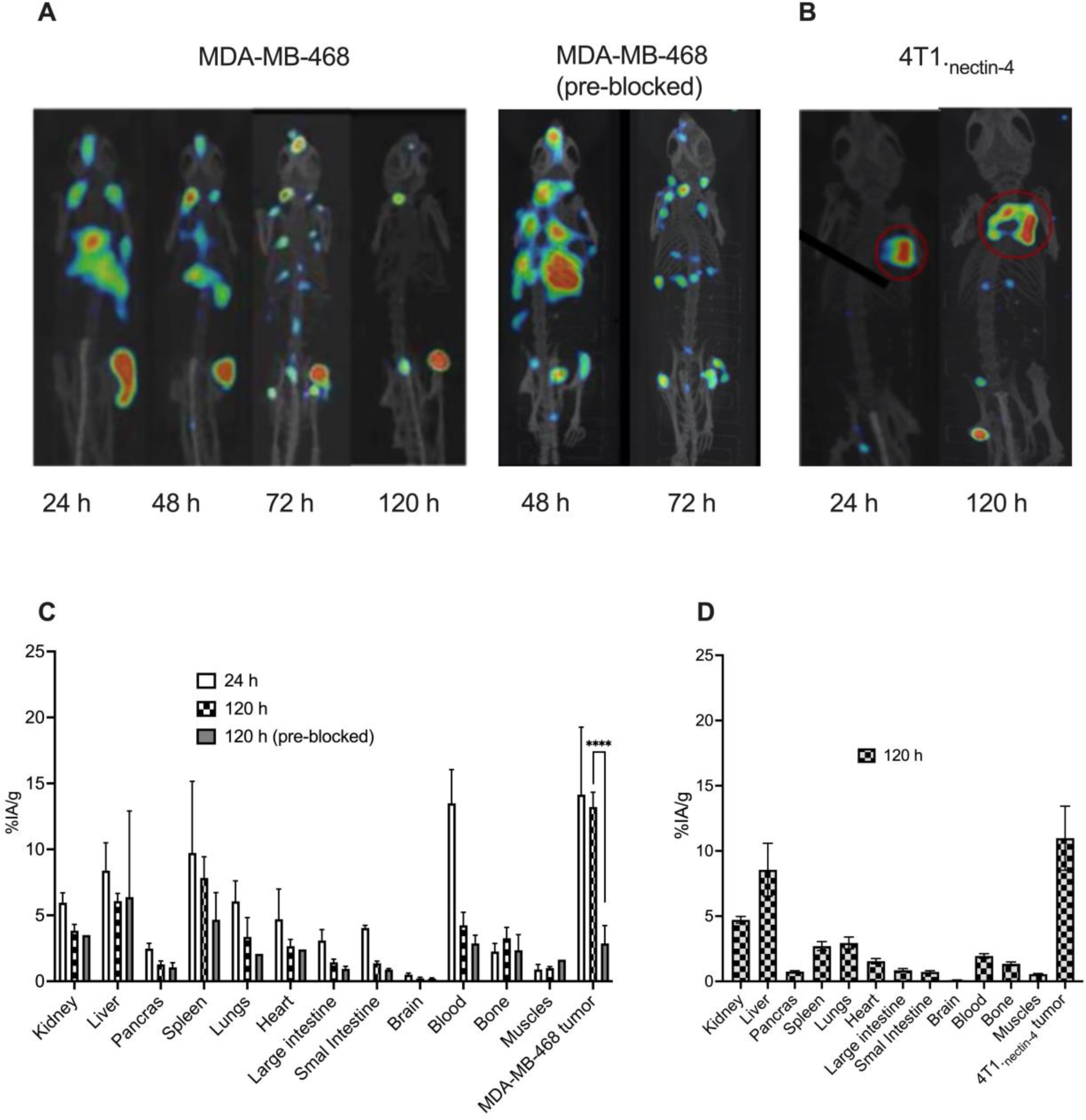
MicroPET/CT imaging and biodistribution of [^89^Zr]Zr-DFO-N4MU01. **A**) Maximum intensity projection (MIP) microPET/CT images of unblocked and N4MU01 (200 µg) pre-blocked CD-1 nude mice bearing nectin-4-positive MDA-MB-468 xenografts (right flank) at different time points p.i. of 12 MBq [^89^Zr]Zr-DFO-N4MU01. **B**) MIP images of the syngeneic mouse model bearing 4T1._nectin-4_ tumors acquired after a 12 MBq [^89^Zr]Zr-DFO-N4MU01 injection. **C**) Biodistribution of [^89^Zr]Zr-DFO-N4MU01 in nectin-4 expressing MDA-MB-468 xenografts and pre-blocked MDA-MB-468 xenografts. **D**) Biodistribution of female BALB/c mice bearing 4T1_nectin-4_ tumors at 120 h p.i.. \*\*\*\**p < 0.0001*

Biodistribution of [^89^Zr]Zr-DFO-N4MU01 was performed at 24 and 120 h p.i. (Figure 3C) in female CD-1 nude mice bearing nectin-4 positive MDA-MB-468 xenograft. To investigate specificity, mice bearing MDA-MB-468 xenografts were pre-blocked using 200 µg of unlabeled N4MU01 4 h prior to injection of [^89^Zr]Zr-DFO-N4MU01. In CD-1 nude mice, tumor uptake of MDA-MB-468 was similar at 24 h (14.1 ± 5.1 % IA/g) and 120 h p.i. (13.2 ± 1.1 %IA/g) (p > 0.9999). Pre-blocking with unlabeled N4MU01 reduced tumor uptake to 2.8 ± 1.3 %IA/g at 120 h (p < 0.0005) p.i. There was a slightly high uptake of [^89^Zr]Zr-DFO-N4MU01 in the lungs (6.0 ± 1.5 % IA/g) and heart (4.7 ± 2.3 % IA/g) at 24 h p.i., likely reflecting blood pool activity. However, this decreased to (3.4 ± 1.5 %IA/g and 2.7 ± 0.5 %IA/g) at 120 h p.i., respectively. Tumor-to-blood ratio for MDA-MB-468 was 1.0 at 24 h and 3.1 at 120 h p.i. The tumour-to-muscle ratio for MDA-MB-468 was 13.2 compared with 1.7 for pre-blocked at 120 h p.i.

Biodistribution of [^89^Zr]Zr-DFO-N4MU01 in mice bearing 4T1_nectin-4_ tumors was performed at 120 h p.i. (Figure 3D) and the tumor uptake was (11 ± 2.5 % IA/g). At 120 h p.i., tumor-to-muscle ratio was 20, and tumor-to-blood ratio was 5.5, which was 0.7 and 0.6-fold higher than observed in the immune-deficient nude mice. Spleen uptake was (2.7 ± 2.5 % IA/g), which was 3.7-fold lower compared to the spleen uptake observed in the immune-deficient nude mice.

### *In vitro* cytotoxicity of [^225^Ac]Ac-Macropa-N4MU01

Live cell imaging was used to study the *in vitro* cytotoxicity of unlabeled N4MU01and [^225^Ac]Ac-Macropa-N4MU01on MDA-MB-468, MCF7 and MDA-MB-231 cells (supplementary Table 1). Phase contrast images showed potent cytotoxicity with [^225^Ac]Ac-Macropa-N4MU01 compared to unlabeled N4MU01. Increased cytotoxicity was observed in MDA-MB-468 at 24 h using [^225^Ac]Ac-Macropa-N4MU01 with IC_50_ of 1.2 kBq/mL, whereas unlabeled N4MU01 had no effect on MDA-MB-468 cells. The IC_50_ values for MCF7 and MDA-MB-231 cells were consistent with moderate and no expression of nectin-4, respectively.

### Biodistribution and dosimetry of [^225^Ac]Ac-Macropa-N4MU01

The biodistribution of [^225^Ac]Ac-Macropa-N4MU01 was studied in healthy BALB/c mice. The uptake of [^225^Ac]Ac-Macropa-N4MU01 was high in the kidney, liver, spleen, lungs, and blood at early time points, but this uptake decreased over time. Ten days p.i., high uptakes were observed in the spleen (4 ± 1.3 %IA/g), lungs (6 ± 1.6 %IA/g), and the blood (5.8 ± 0.9 %IA/g) (Supplementary Table 2). Human organ dosimetry estimates of [^225^Ac]Ac-Macropa-N4MU01 (Table 1) show the organs receiving the highest dose are the lungs>liver>spleen and the total whole-body dose estimate of 3.59 mSv/MBq.

**Table 1.** Projected human absorbed doses (mSv/MBq) of [^225^Ac]Ac-Macropa-N4MU01.

| **Organs** | **mSv/MBq female** |
| --- | --- |
| Adrenals | 6.20E-03 |
| Brain | 8.50E-01 |
| Breast | 7.66E-04 |
| Esophagus | 3.16E-03 |
| Eyes | 8.08E-05 |
| Gall bladder | 2.69E-03 |
| Left colon | 1.01E-03 |
| Small intestine | 7.21E-04 |
| Stomach | 2.15E-03 |
| Right colon | 8.42E-04 |
| Rectum | 1.22E-04 |
| Heart wall | 1.20E01 |
| Kidneys | 5.59E01 |
| Liver | 6.21E01 |
| Lungs | 9.04E01 |
| Ovaries | 1.63E-04 |
| Pancreas | 1.42E01 |
| Salivary glands | 1.67E-04 |
| Red marrow | 9.35E-04 |
| Spleen | 7.65E01 |
| Thymus | 3.18E-03 |
| Thyroid | 1.15E-03 |
| Bladder | 1.82E-03 |
| Uterus | 1.54E-04 |
| Total body | 3.59E00 |
| Effective dose | 1.49E01 |

### Efficacy of [^225^Ac]Ac-Macropa-N4MU01

We studied the efficacy of [^225^Ac]Ac-Macropa-N4MU01 in a TNBC MDA-MB-468 mouse xenograft. Animals treated with two doses of 13 kBq [^225^Ac]Ac-Macropa-N4MU01 or 18.6 kBq [^225^Ac]Ac-Macropa-N4MU01 had a remarkable decrease in tumor volume compared to saline and unlabeled N4MU01 controls (Figure 4A, individual mouse tumor growth curves are shown in Supplementary Figure 5). All the mice in saline group reached the study endpoint of ≥ 1500 mm^3^ by day 21. One of 6 mice (16.6%) treated with unlabeled N4MU01 reached the study endpoint by day 21, and the rest reached the endpoint by day 24. The Kaplan-Meier survival curve showed that mice treated with [^225^Ac]Ac-Macropa-N4MU01 survived for the whole period of the therapy study of 30 days. However, the median survival of the saline group was 21 days, and the median survival of the unlabeled N4MU01-treated mice was 23.5 days (Figure 4B).

**Figure 4:**
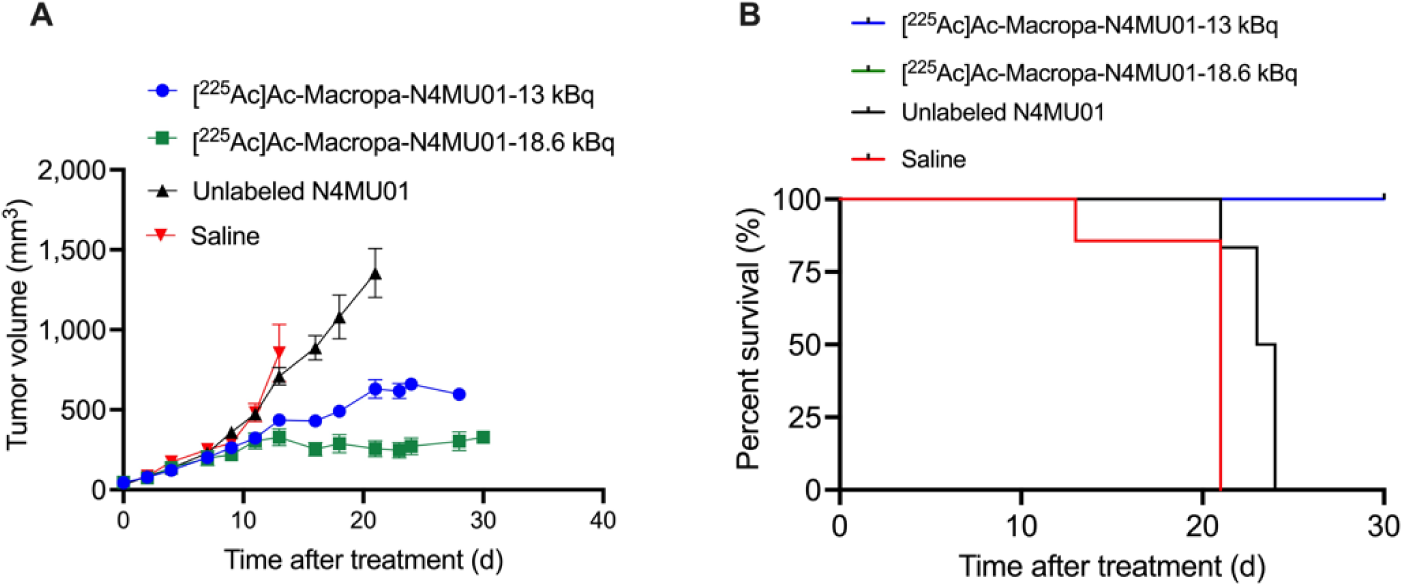
Efficacy of [^225^Ac]Ac-Macropa-N4MU01 in MDA-MB-468 xenografts mouse model. **A**) Average tumor volumes of MDA-MB-468 xenografts (n = 4-7 mice/group). **D**) Kaplan-Meier survival curve of mice bearing MDA-MB-468 xenograft. Study endpoint was considered as tumor volume ≥ 1500 mm^3^.

To further assess the efficacy of [^225^Ac]Ac-Macropa-N4MU01, a 4T1._nectin-4_ syngeneic BALB/c mouse model was used (Figure 5A). The transfection of mouse breast cancer cell line 4T1 with human nectin-4 was confirmed using flow cytometry as shown in (Supplementary Figure 6). Mixed population of nectin-4 transfected and non-transfected 4T1 cells was observed. Cell sorting using FACS analysis was performed to isolate the nectin-4 transfected 4T1 population (Supplementary Figure 7).

**Figure 5:**
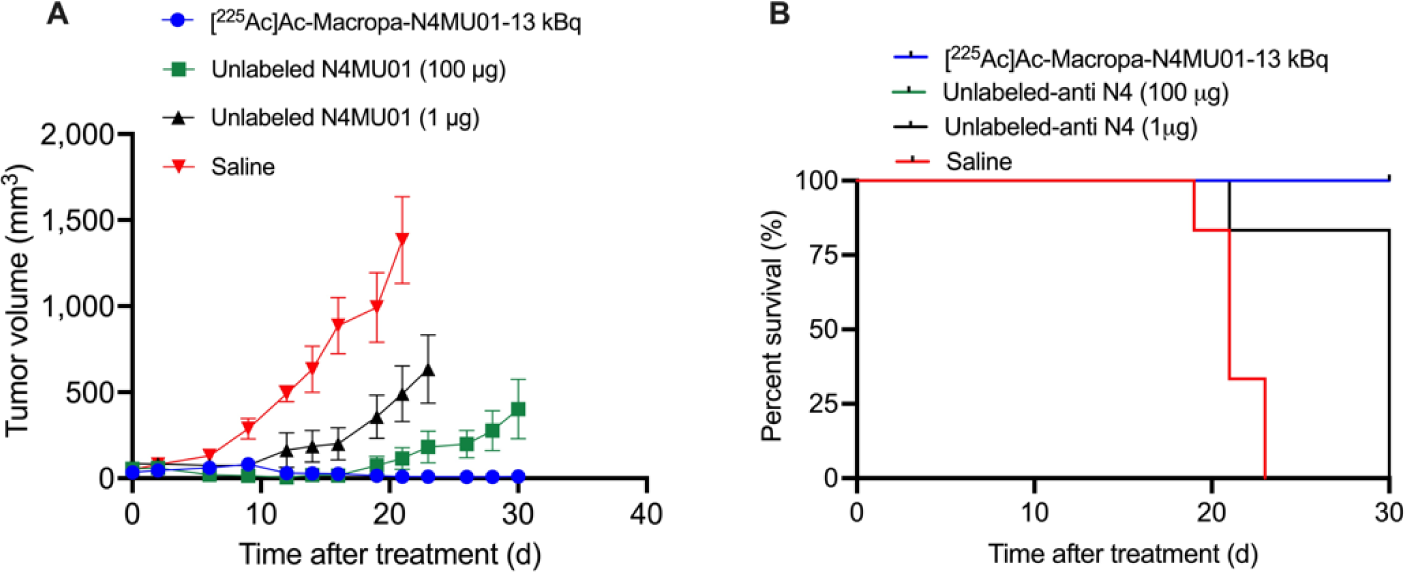
Efficacy of [^225^Ac]Ac-Macropa-N4MU01 in 4T1._nectin-4_ syngeneic mouse model. **A**) Average tumor volumes (n = 5-6 mice/group). **B**) Kaplan-Meier survival curve. Study endpoint was considered as tumor volume ≥ 1500 mm^3^.

For the syngeneic mouse model, five of six mice (83.4%) treated with two doses of 13 kBq [^225^Ac]Ac-Macropa-N4MU01 had complete remission by day 30. Five of six mice (83.4%) treated with the therapeutic dose of unlabeled N4MU01 (100 μg) had delayed tumor growth, while 1/6 mice (16.6%) had complete remission (Figure 5A, tumor growth curves of individual mice are shown in Supplementary Figure 8). All the mice in saline and the unlabeled N4MU01 groups reached the study endpoint of ≥ 1500 mm^3^ by day 23 and 30, respectively. The median survival for mice treated with [^225^Ac]Ac-Macropa-N4MU01 and unlabeled N4MU01 therapeutic dose (100 μg) was not reached by day 30. However, the median survival for the saline was 21 days (Figure 5B). There was no apparent toxicity throughout the treatment period, evident from the weight gain (Supplementary Figure 9A and B).

## Discussion

With the exception of sacituzumab govitecan, there are no other targeted therapies available for TNBC, and surgery followed by chemotherapy remains the current standard of care [26]. An optimal cancer cell surface biomarker is essential for developing efficient targeted therapeutics against TNBC. Nectin-4 is overexpressed in ∼62% of TNBCs and in TNBC metastases and basal subtypes and absent in normal epithelial breast tissue [6] making it an ideal target for theranostics. For clinical application, fully human antibodies are advantageous for therapy for which the concern of immunogenicity is eliminated [27]. This study describes the preclinical evaluation of a fully human anti-nectin-4 antibody as a PET imaging probe for the early detection of TNBCs and its development as a radioimmunotherapeutic agent. While a few groups have evaluated anti-nectin-4 immunoPET/SPECT imaging, photothermal probes and ADCs, no reports of alpha particle (^225^Ac) radioimmunotherapy (RIT) of nectin-4 exist in the literature [28, 29]

^89^Zr is considered an ideal PET isotope because of its long half-life of 78.4 h and high resolution due to the emission of positrons [30]. To evaluate N4MU01 IgG as immunoPET imaging probe, anti-nectin-4 antibody was conjugated with *p*-SCN-Bn-DFO, and the immunoconjugate had similar low nanomolar affinity (3 nM) compared with the unconjugated antibody. As expected, radiolabeling with ^89^Zr resulted in a stable probe. The imaging [^89^Zr]Zr-N4MU01 probe was evaluated (microPET and biodistribution studies) using nectin-4 positive TNBC xenografts with high and medium receptor densities. Shao *et al* [29] studied the tumor uptake of [^99m^Tc]Tc-HYNIC-mAb_Nectin-4_ in nectin-4 positive MDA-MB-468 xenograft. Maximum tumor uptake in MDA-MB-468 xenograft was 15.32 ± 1.04% ID/g and when pre-blocked was 4.33 ± 0.48% ID/g which is similar with our study. Additionally, the tumor-to-blood and tumor-to-muscle ratios for our study are comparable with those for [^99m^Tc]Tc-HYNIC-mAb_Nectin-4_ tracer. Campbell *et al* evaluated ^89^Zr-AGS-22M6 in tumor bearing mice and cynomolgus monkeys [28]. Tumor uptake of [^89^Zr]Zr-AGS-22M6 in nectin-4 transduced MDA-MB-231 cells (MDA-MB-231-nectin-4) transduced to express the receptor ranged from an average of 38.8 ± 2.8 % ID/g on day 1 to an average of 39.9 ± 5.9 %ID/g on day 6 with an average high of 45.3 ± 2.4 %ID/g on day 3 compared with non-transduced receptor negative MDA-MB-231-neo) that ranged from ranged from an average of 16.3 ± 1.8 %ID/g on day 1 to an average of 17.2 ± 1.3 %ID/g on day 6 with an average high of 18.2 ± 2.8 % ID/g on day 2. Similar tumor uptake ratios were found for [^89^Zr]Zr-AGS-22M6 in nectin-4 positive patient-derived xenografts (PDX) compared with nectin-4 negative PDXs. In the current study, tumor uptake of [^89^Zr]Zr-DFO-N4MU01 in MDA-MB-468 decreased from 13.2 %IA/g to 2.8 %IA/g when pre-blocked with cold antibody which is indicative for better specificity compared with [^89^Zr]Zr-DFO-AGS-22M6 in spite of its apparent lower K_D_ of 0.01 nM [8]. In addition, [^89^Zr]Zr-DFO-N4MU01displayed fast distribution half-life t_1/2α_ of 3.37 h and a relatively moderate clearance t_1/2β_ of 63 h compared with other very slow-clearing IgGs such as trastuzumab which is advantageous for imaging.

Eighteen-membered ring macrocylic chelator Macropa forms a highly stable and inert complex with actinium-225 [22, 23]. To investigate the antitumor effects of the radioimmunoconjugate, we studied the *in vitro* cytotoxicity using IncuCyte S3 live-cell imaging with Cytotox Red reagent, that allows for real-time quantification of dead cells. [^225^Ac]Ac-Macropa-N4MU01 displayed enhanced cytotoxicity to nectin-4 expressing cells while unlabeled N4MU01 was not cytotoxic to nectin-4 expressing TNBC MDA-MB-468. The IC_50_ of (1.2 kBq/mL) was similar to the other highly cytotoxic [^225^Ac]Ac-Macropa-labeled radioimmunoconjugates [31, 32].

Our [^225^Ac]Ac-Macropa-N4MU01 showed very favorable dosimetry in all healthy organs due to its excellent clearance rates from nectin-4 negative healthy tissues. The highest organ dose was observed for the liver, lungs and spleen. No organ dose estimates for anti-nectin-4 radioimmunoconjugate (RIC) has been reported in the literature for comparison. By comparison widely investigated anti-HER2 RIC such as [^177^Lu]Lu-DTPA-trastuzumab has liver and spleen doses of 1.72 Gy/MBq (1720 mSv/MBq) and 1.6 Gy/MBq (1600 mSv/MBq), respectively which is several folds higher than [^225^Ac]Ac-Macropa-N4MU01 [33]. [^225^Ac]Ac-Macropa-N4MU01 (total body dose of 3.59 mSv/MBq) was almost 8-fold less than the widely investigated peptide targeted radioconjugate [^225^Ac]Ac-DOTA-PSMA-617 (28 mSv/MBq) [34].

We then investigated the therapeutic efficacy of the unlabeled N4MU01 and [^225^Ac]Ac-Macropa-N4MU01 at controlling the growth of the TNBC MDA-MB-468 xenografts. Mice were administered two doses of 13 kBq (low dose) or 18.6 kBq (high dose) on days 0, and 10. Although we have not studied the safety and maximum tolerated dose of [^225^Ac]Ac-Macropa-N4MU01 in mice, the low and high doses were picked based on our experience with other antigens/radioimmunoconjugates. All saline treated mice bearing MDA-MB-486 xenograft reached tumor endpoint of 1500 mm^3^ by day 21. Significant dose dependent tumor inhibition was for the 13 kBq and 18.6 kBq doses at day 28 compared with controls. Additionally, we evaluated the effectiveness of [^225^Ac]Ac-Macropa-N4MU01 in a syngeneic mouse model after transfection of murine 4T1 cells with nectin-4 (4T1_nectin-4_). 5/6 mice bearing 4T1_nectin-4_ treated using two doses of 13 kBq [^225^Ac]Ac-Macropa-N4MU01 had complete remission while the remaining mouse had a tumor volume of 8.2 ±14.0 mm^3^ at day 28. Untreated mice and mice treated using 1 µg (the equivalent antibody mass dose of the ^225^Ac-labeled agent) reached the tumor volume of >1500 mm^3^ by day 21. Unlike the ADC studies reported by M-Rabet *et al*. [7] and Challita-Eid *et al.* [8], tumor re-growths were not observed using [^225^Ac]Ac-Macropa-N4MU01.

## Conclusion

In this study, we describe the preclinical evaluation of a fully human anti-nectin-4 antibody for the non-invasive PET imaging and the targeted RIT of TNBC. Here, we report the development of the first [^225^Ac]Ac-Macropa-N4MU01 α-particle therapeutic agent. Our N4MU01 antibody is fully human and, therefore, is less immunogenic compared with murine-based antibodies, which is essential for translation into effective pharmaceuticals. The targeting specificity, the high tumor uptake of [^89^Zr]Zr-DFO-N4MU01, and the exceptional therapeutic efficacy of [^225^Ac]Ac-Macropa-labeled N4MU01 warrants further evaluation in different mouse models and potential clinical translation.

## Ethics Statement

Animal Studies: All animal studies were approved by the University of Saskatchewan Animal Care and Use Committee protocol #20220021.

## Author contributions

H.F. and M.U. conceived, designed, supervised the study and edited the manuscript, H.B drafted the initial version of the manuscript, H.B and F.N.N. performed experimental design, execution, and data analysis, with technical assistance from J.P.K., A.F.T., A.D., and EN. All authors have read and agreed to the final version of the manuscript.

## Supplementary material

Supplementary figures 1-8 and figure legends are described in attached document.

