## Supplementary Information for "[^225^Ac]Ac/[^89^Zr]Zr-labeled N4MU01 radioimmunoconjugates as theranostics against nectin-4 positive triple negative breast cancer"

### Supplementary Methods

#### Cell lines and xenografts

Human breast cancer cell lines MDA-MB-468, MCF-7 which expresses nectin-4, and MDA-MB-231, known for nectin-4 negative expression, were purchased from ATCC (Rockville, MD). MDA-MB-468 and MDA-MB-231 cells were propagated in Leibovitz's L-15 medium (HyClone Laboratories, Logan, UT, USA) supplemented with 10% FBS. MDA-MB-468 and MDA-MB-231 cells were incubated at 37 °C without $CO_2$. MCF-7 cells were cultured in DMEM mediam (HyClone Laboratories, Logan, UT, USA) supplemented with 10% FBS. For syngeneic mouse models, mouse breast cancer cell line 4T1 was purchased from ATCC (Rockville, MD) and transfected with full-length nectin-4 protein ($4T1._{nectin\text{-}4}$). $4T1._{nectin\text{-}4}$ cells were propagated in RPMI 1640 medium (HyClone Laboratories, Logan, UT, USA), supplemented with 10 % FBS and incubated at 37 °C in a humidified atmosphere of 5 % $CO_2$.

Female CD-1 nude, athymic nude, and immune-competent BALB/c mice of 4 weeks of age were obtained from Charles River, Canada (St-Constant, QC, Canada). Animals used in this study were maintained following the guidelines of the University of Saskatchewan Animal Care Committee (protocol # 20170084 and 20220021). At five weeks of age, CD-1 nude mice were subcutaneously injected at the right flank with $5 \times 10^6$ MDA-MB-468 cells in 100 μL suspension of a 1:1 mixture of medium with matigel (Discovery Laboware, Inc. Bedford, MA). BALB/c mice were subcutaneously injected at the right flank with $2.5 \times 10^6$ 4T1 cells transfected with human nectin-4 in 100 μL PBS. Tumor growth was measured using a digital caliper.

#### 4T1 cell transfection with human nectin-4

Human Nectin-4 (NP_112178) Versa Clone cDNA (Cat # RDC1544, &D Systems, Inc.) (accession # NP_112178) was purchased from R&D systems, Inc. The coding sequence of nectin-4, excluding the signal peptide (aa 32-510) was amplified by PCR using the following primers;

FN19 5‘-
CTAAGTCTTGCACTTGTCACGAATTCGATATCGCAGGGTGAGCTGGAGACCTCAG-3’

FN20 5‘-
CTTAACGCGCCACCGGTTAGCGCTAGCATGCATACTCAGACCAGGTGTCCCCGC-3’

The PCR product was cloned into a custom lentiviral vector pMUFV01 linearized after digestion with enzymes (EcoRI and NheI) using Gibson assembly. The fully assembled vector map is shown in Supplementary Figure 1.

Sequence verified plasmid was transfected into 4T1 mouse breast cancer cell line. Briefly, 5 x $10^5$ 4T1 cells in 3 mL of growth medium were seeded in a 6-well culture plate 24h before transfection. On the day of transfection, the seeding medium was removed, and cells were washed with pre-warmed DPBS. The transfection medium was prepared by mixing 2 mg of plasmid DNA, 100 mL of growth medium, and 10 mL of FuGENE HD transfection reagent (Cat # E2311, Promega) and incubating for 15 min. The incubated mixture was then added to washed cells, and the volume was made up to 3 mL with a complete growth medium. The plate was then incubated at 37 °C in a humidified atmosphere of 5 % $CO_2$. 24 h following transfection, cells were trypsinized, washed and seeded in a growth medium supplemented with 2mg/mL Blasticidin.

**Flow cytometry and sorting of human nectin-4 transfected cells**

Nectin-4 cDNA transfected 4T1 cell pool was sorted for high nectin-4 expression using flow cytometry. Cells were labeled with 500 nM N4MU01 followed by secondary antibody Goat anti-Human IgG PE-conjugated secondary antibody (eBioscience, cat. #12-4998-82). Sorting was performed using the BD FACS Melody cell sorter (BD Biosciences) to select high nectin-4 overexpressing population of $4T1._{nectin\text{-}4}$. The sorted cells were then propagated for making stocks and expanded for injection into mice.

**Characterization of N4MU01 binding using flow cytometry**

In a 96 well non-binding plate, different concentrations of the N4MU01 in 1 x PBS of (500 nM, 250 nM, 125 nM, 62 nM, 30 nM, 10 nM and 3 nM) were added to nectin-4 expressing breast cancer cells MDA-MB-468, MCF-7 and nectin-4 negative MDA-MB-231 cells at 3 x $10^5$ cells/well. After 30 min of binding at 4 °C, cells were washed two times and re-suspended in ice-cold 1x PBS. FITC labeled goat F(ab')2 fragment anti-human IgG (H + L) antibody (Beckman Coulter, Brea, CA, USA) was added to cells in (1 in 50) dilution and allowed to bind for 30 min at 4 °C. Cells were washed with ice-cold 1 x PBS three times before reading the plate using CytoFLEX (Beckman Coulter) on the FL1 channel. Flow cytometry data were acquired using

CytoFLEX and analyzed using FlowJo (version 10.7.2; FlowJo LLC, Ashland, OR, USA). GraphPad Prism v9 (GraphPad software, San Diego, CA, USA) was used to determine the binding constant ($K_d$) and half maximal effective concentration ($EC_{50}$) [1].

**Internalization of N4MU01**

In a flat-bottom 96-well plate, nectin-4 expressing MDA-MB-468, MCF-7 and nectin-4 negative MDA-MB-231 cells were seeded and incubated at 37 °C overnight. Twenty-four hours later, anti-nectin-4 antibody with a final concentration of 4 µg/mL was added to a 3x molar-excess of FabFluor pH red antibody internalization reagent (Essen Bioscience Ann Arbor, MI, USA) and incubated for 15 minutes at 37 °C. The labeled antibody was then added to the cells, and images were taken at 10x magnification every 2 hours with phase contrast and red fluorescence filters using the Incucyte live-cell imaging system (Essen BioScience Ann Arbor, MI, USA). The mean red object area ($\mu m^2$/well) was calculated using the Incucyte software and was used to quantify antibody internalization.

**Conjugation with bifunctional chelators DFO and Macropa and radiolabeling with [$^{89}$Zr]Zr and [$^{225}$Ac]Ac**

The conjugation of N4MU01 to desferrioxamine (DFO) was carried out using lab SOPs [2]. To conjugate the N4MU01 with the macropa chelator, a stock of 20 mg/mL of macropa-SCN was prepared for conjugation reactions. Briefly, 1 mg each of the above antibodies was buffer exchanged with 0.1 M sodium bicarbonate buffer containing 0.15 M sodium chloride using a spin cap column. After buffer exchange, the desired volume of macropa-SCN (15-fold excess) was added to the antibody solution and incubated at 4˚C for 18 h. Following incubation, excess unreacted macropa was removed using Amicon Ultra-4 10K centrifugal filters (EMD Millipore, Burlington, MA, USA). Radiolabeling of DFO-N4MU01 and Macropa-N4MU01 with [$^{89}$Zr]Zr-oxalate and [$^{225}$Ac]Ac-nitrate was carried out as reported earlier [2-4]

**Radioligand binding assay of [$^{89}$Zr]Zr-DFO-N4MU01 and [$^{225}$Ac]Ac–Macropa-N4MU01**

The binding of [$^{89}$Zr]Zr-DFO-N4MU01 to nectin-4 positive MCF-7 cells and [$^{225}$Ac]-Ac-macropa-N4MU01 to nectin-4 expressing MDA-MB-468 was determined using a saturation radioligand binding assay. Briefly, $10^6$ cells were incubated with increasing concentrations of [$^{89}$Zr]Zr-DFO-

N4MU01 and [$^{225}$Ac]Ac-Macropa-N4MU01 (0.17 to 375 nmol/L and 0.19 to 12.5 nmol/L in 100 µL of PBS, respectively) for 4 h at 4°C. Non-specific binding was determined in a similar assay using a 10-fold molar excess of unlabeled anti-nectin-4 antibody for binding of [$^{89}$Zr]Zr-DFO-anti-nectin-4 antibody and 50 molar excess for [$^{225}$Ac]Ac-Macropa-N4MU01. A non-linear regression analysis with one-site - total and non-specific binding equation was used to determine the $K_D$ using GraphPad Prism version 9 (La Jolla, CA, USA).

**MicroPET/CT imaging and biodistribution of N4MU01 radioimmunoconjugates**

Female CD-1 nude mice (n = 4/group) bearing nectin-4 positive MDA-MB-468 xenograft and female BALB/c athymic nude mice (n = 4) bearing nectin-4 positive BT-474 and MCF-7 xenografts were injected intravenously using 12 ± 1 MBq (21 – 26 µg) [$^{89}$Zr]Zr-DFO-N4MU01. Additionally, mice bearing MDA-MB-468 xenograft (n = 4) were injected intravenously with 200ug of unlabeled N4MU01 four hours prior to the injection of [$^{89}$Zr]Zr-DFO-N4MU01 to pre-block nectin-4 receptors. For the syngeneic mouse model, female BALB/c mice (n=4) bearing $4T1_{nectin\text{-}4}$ tumors were injected intravenously with 12 ± 1 MBq (21 – 26 µg) [$^{89}$Zr]Zr-DFO-N4MU01. MicroPET/CT imaging was performed at 24, 48, 72, 96, and 120 h post-injection and at 24 and 120 h for the syngeneic mouse model using the Vector$^4$CT scanner (MILabs B.V., Utrecht). PET scans were acquired in a list-mode data format using a high-energy ultra-high resolution (HE-UHR-1.0 mm) mouse/rat pinhole collimator. Images were reconstructed using PMOD 3.8 software (PMOD, Switzerland). Immediately after imaging, mice were sacrificed at 120 h post injection for biodistribution studies. To complete the biodistribution studies, an additional group (n = 4) of MDA-MB-468 tumor-bearing mice were sacrificed 24 h post injection. All major organs and blood were harvested, collected in tared tubes and weighed. The radioactivity in all organs and blood was measured using an automated gamma counter (Wallac Wizard 1480, PerkinElmer, Waltham, MA), and expressed as percent injected activity per gram (%IA/g).

**Pharmacokinetic profile of [$^{89}$Zr]Zr-DFO-N4MU01**

Female CD-1 nude mice (n = 4) were injected intravenously with 5 – 6 MBq of [$^{89}$Zr]Zr-DFO-N4MU01 (10 – 12 µg of antibody). Blood was collected from a saphenous vein in heparinized capillary tubes at different time points (0.5 – 120 h). The height of the capillary tube occupied by blood was determined using a digital caliper. Blood volume in the capillary tube (mL) was measured using $V = \pi\ r^2 h$. The radioactivity in the samples was measured using a gamma counter and expressed as % injected activity/mL (%IA/mL). Pharmacokinetic parameters, including the distribution and elimination half-lives ($t_{1/2\alpha}$ and $t_{1/2\beta}$), volume of distribution at steady-state (Vss), clearance (CL) and volume of the central compartment ($V_1$), were calculated by fitting the blood radioactivity versus time curve to a two-compartment model with i.v. bolus input.

### *In vitro* cytotoxicity

Live-cell images were captured every 2 h using a 10Å~ objective lens using phase contrast and fluorescence channel. Images were processed and analyzed using IncuCyte S3 software. The red fluorescent values were generated, and $IC_{50}$ values for individual compounds were calculated using GraphPad prism v9.

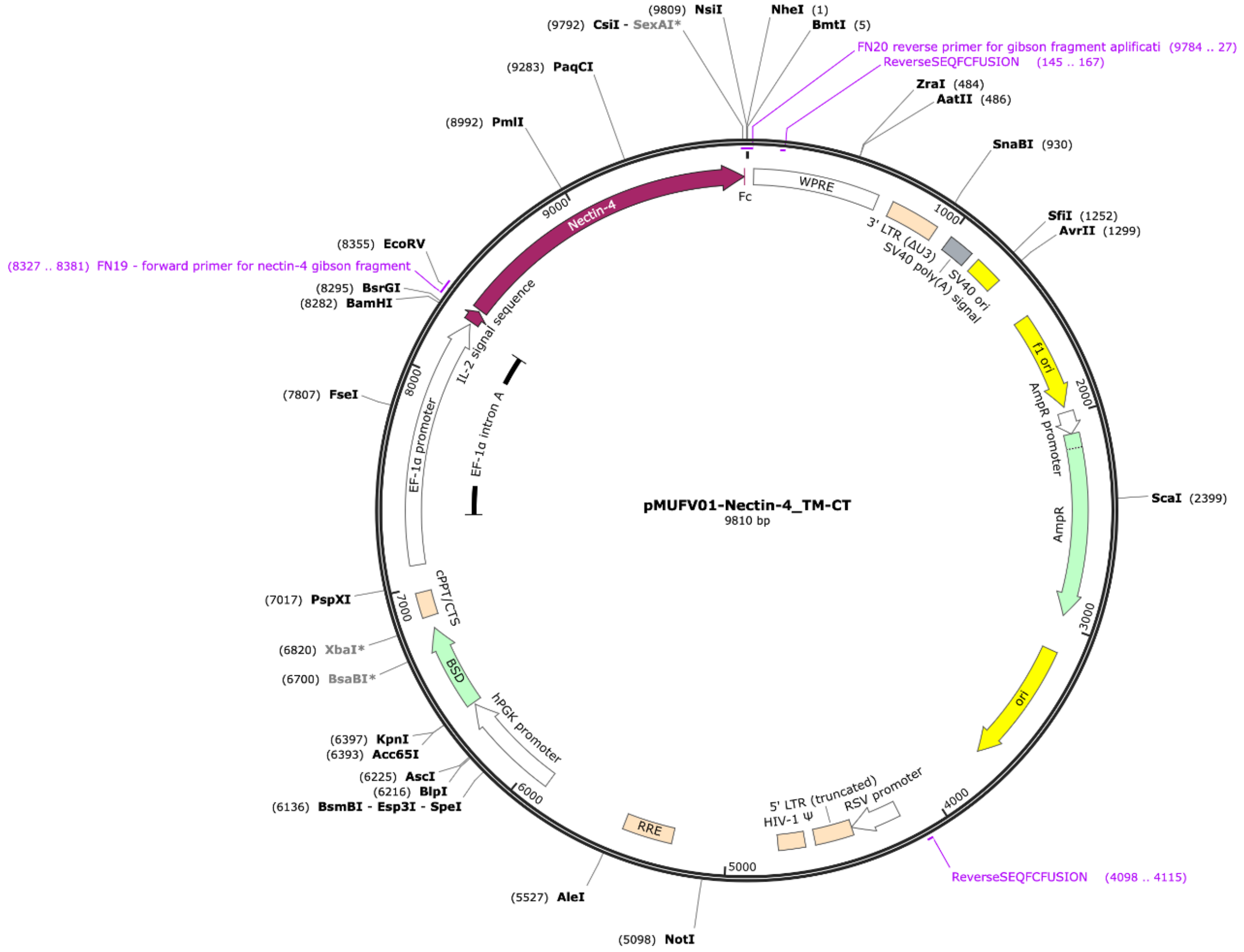

**Supplementary Figure 1:** pMUFV01-Nectin-4, a custom lentiviral vector used to clone hNectin-4 cDNA.

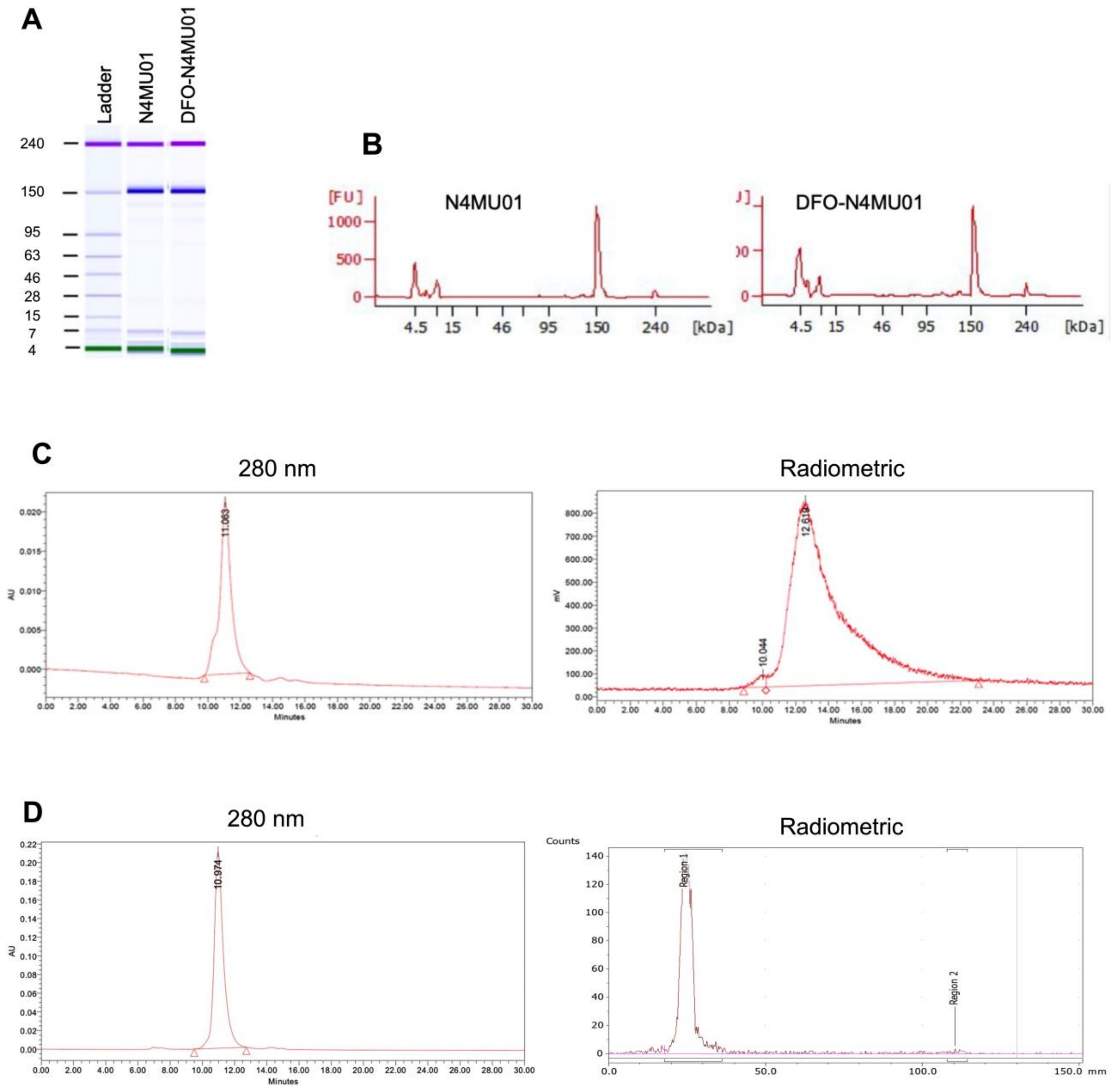


**Supplementary Figure 2**: Bioanalyzer ladder (A) and chromatograms (B) of unconjugated N4MU01, and DFO-N4MU01. C) Representative size exclusion (SEC) HPLC chromatograms showing purity of DFO-N4MU01 and [$^{89}$Zr]Zr-DFO-N4MU01. UV channel (280) for DFO-N4MU01 and radiometric channels for [$^{89}$Zr]Zr-DFO-N4MU01 are shown. D) Representative size exclusion (SEC) HPLC chromatograms showing purity of Macropa-N4MU01 (280 nm) and iTLC for the high radiochemical yield of [$^{225}$Ac]Ac-Macropa-N4MU01 (radiometric).

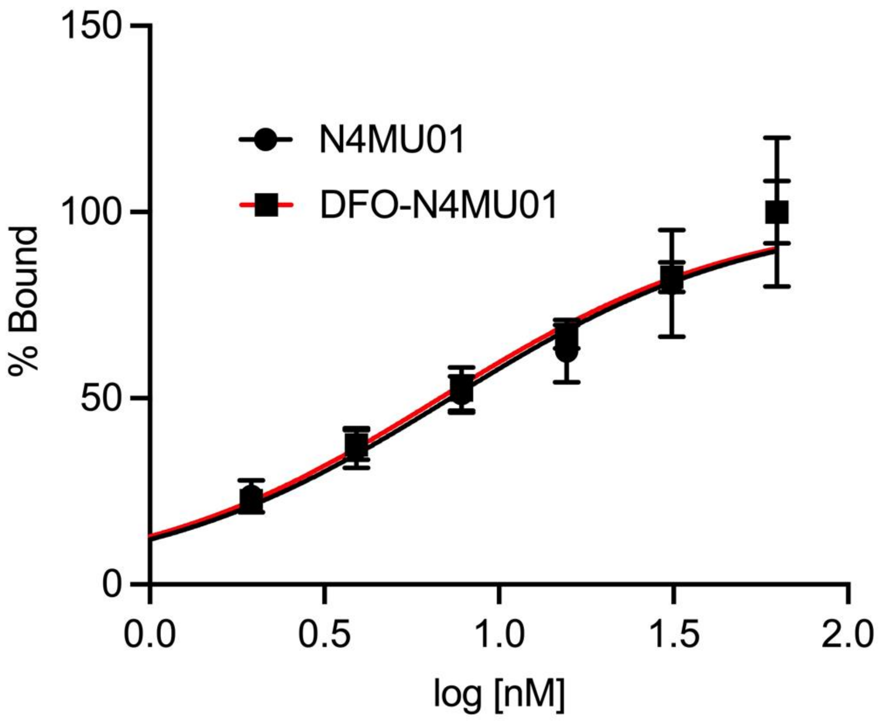


**Supplementary Figure 3:** Estimation of $EC_{50}$ values using saturation binding assay using flow cytometry: Nectin-4 positive MCF-7 cells were titrated with decreasing concentrations of N4MU01 and DFO-N4MU01. Mean fluorescence intensity (MFI) was plotted against concentration.

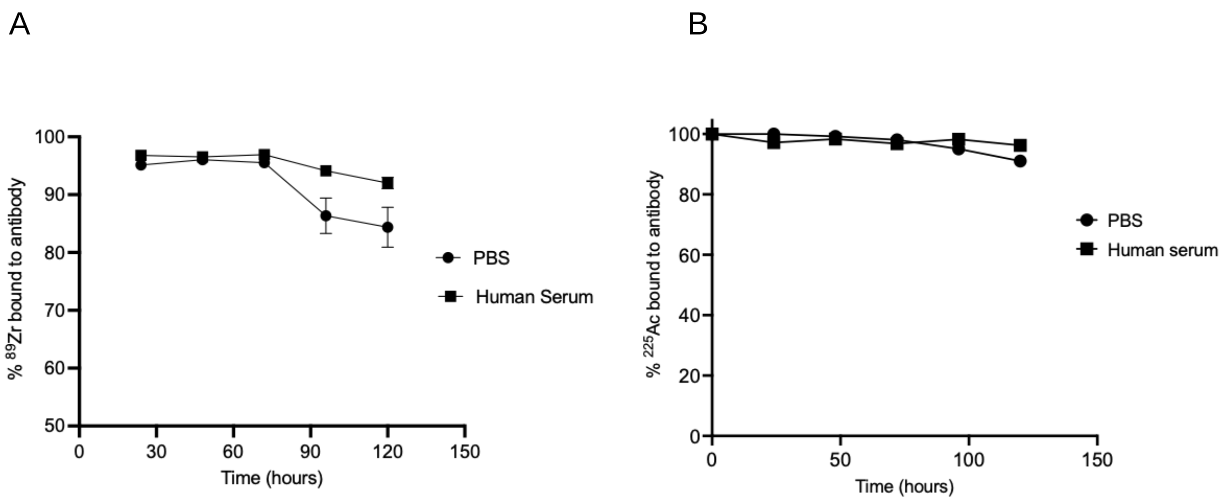


**Supplementary Figure 4**: Stability studies of (A) [$^{89}$Zr]Zr-DFO-N4MU01 and (B) [$^{225}$Ac]Ac-Macropa-N4MU01 in human plasma and PBS at 37°C.

**Supplementary Table 1**: $IC_{50}$ values of immunoconjugates in different nectin-4 expressing TNBC cell lines.

| $IC_{50}$ | MDA-MB-468 | MCF-7 | MDA-MB-231 |
|---|---|---|---|
| N4MU01 (nM) | - | - | - |
| [$^{225}$Ac]Ac-Macropa-N4MU01 (nM) | 0.1 ± 0.05 | 42.6 ± 0.0 | 66600 |
| [$^{225}$Ac]Ac-Macropa-N4MU01 (kBq/mL) | 1.2 ± 0.57 | 64.1 ± 0.0 | $10^5$ |
| [$^{225}$Ac]Ac-Macropa -rituximab (kBq/mL) | 202.7 | 22.5 | 78.5 |

**Supplementary Table 2:** Biodistribution of [$^{225}$Ac]Ac-macropa-N4MU01 in healthy BALB/c mice at different time points post injection expressed as % injected activity per gram (%IA/g) ± SD

| Organs | 1 h | 24 h | 120 h | 240 h |
|---|---|---|---|---|
| Bladder | 0.18± 0.22 | 4.43± 2.32 | 4.1± 2.27 | 6.28± 3.33 |
| Kidney | 10.07± 6.08 | 8.8± 2.05 | 4.23± 1.62 | 3.79± 1.16 |
| Liver | 8.79± 3.53 | 6.44± 1.59 | 3.22± 1.08 | 3.8± 0.98 |
| Pancras | 0.56± 0.51 | 2.34± 0.32 | 1.03± 0.49 | 0.77± 0.6 |
| Spleen | 13.37± 6.62 | 9.45± 1.56 | 4.51± 2.09 | 3.81± 1.33 |
| Lungs | 12.75± 6.93 | 12.39± 3.26 | 6.54± 2.72 | 5.91± 1.64 |
| Heart | 6.1± 3.59 | 7.87± 2.12 | 2.57± 0.76 | 2.28± 0.24 |
| Large intestine | 0.85± 0.41 | 2.83± 0.59 | 1.28± 0.68 | 1.18± 0.64 |
| Small Intestine | 2.5± 1.22 | 2.97± 0.91 | 1.34± 0.5 | 1.27± 0.32 |
| Stomach | 0.53± 0.34 | 2.47± 0.52 | 0.85± 0.12 | 1.32± 0.45 |
| Skull | 1.86± 0.94 | 4.31± 0.79 | 2.34± 1.01 | 2.26± 0.41 |
| Brain | 0.33± 0.16 | 0.3± 0.05 | 0.12± 0.12 | 0.05± 0.07 |
| Spine | 2.43± 1.44 | 3.08± 0.64 | 1.29± 0.54 | 1.46± 0.14 |
| Blood | 33.39± 8.64 | 18± 6.94 | 5.5± 2.36 | 5.14± 0.92 |
| Bone | 2.09± 0.97 | 3.09± 1.11 | 1.57± 0.44 | 1.45± 0.64 |
| Muscles | 0.37± 0.4 | 1.62± 0.65 | 0.84± 0.52 | 0.84± 0.46 |
| skin | 1.12± 0.54 | 7.07± 2.2 | 3.73± 0.92 | 3.28± 0.64 |

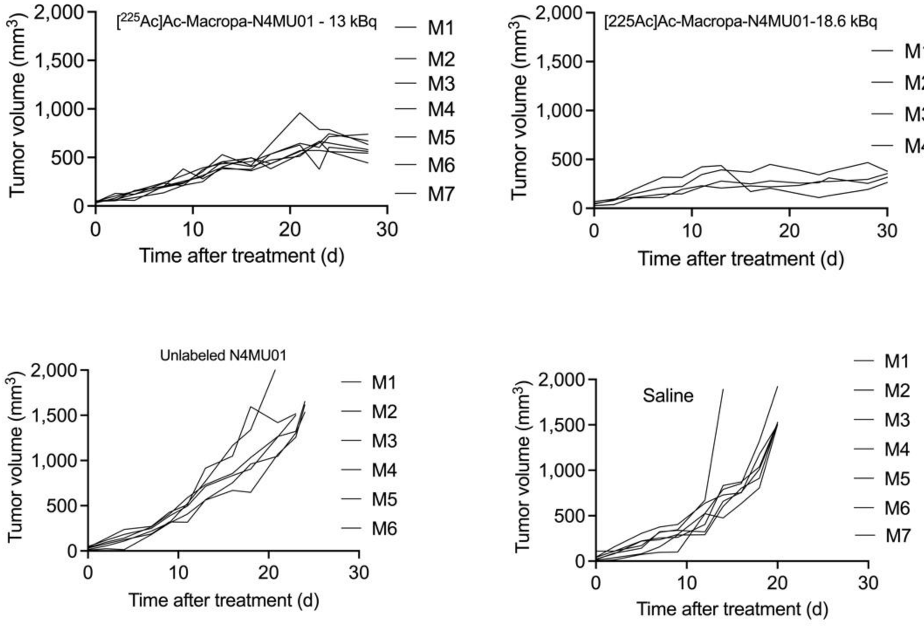


**Supplementary Figure 5:** Efficacy of [$^{225}$Ac]Ac-macropa-N4MU01 radioimmunoconjugate in MDA-MB-468 xenografts model. (A) Tumor growth curve of 13 kBq [$^{225}$Ac]Ac-macropa-N4MU01 18.6 kBq [$^{225}$Ac]Ac-macropa-N4MU01, saline control and unlabeled N4MU01-treated mice bearing MDA-MB-468 xenograft represented as tumor volumes versus days post-treatment.

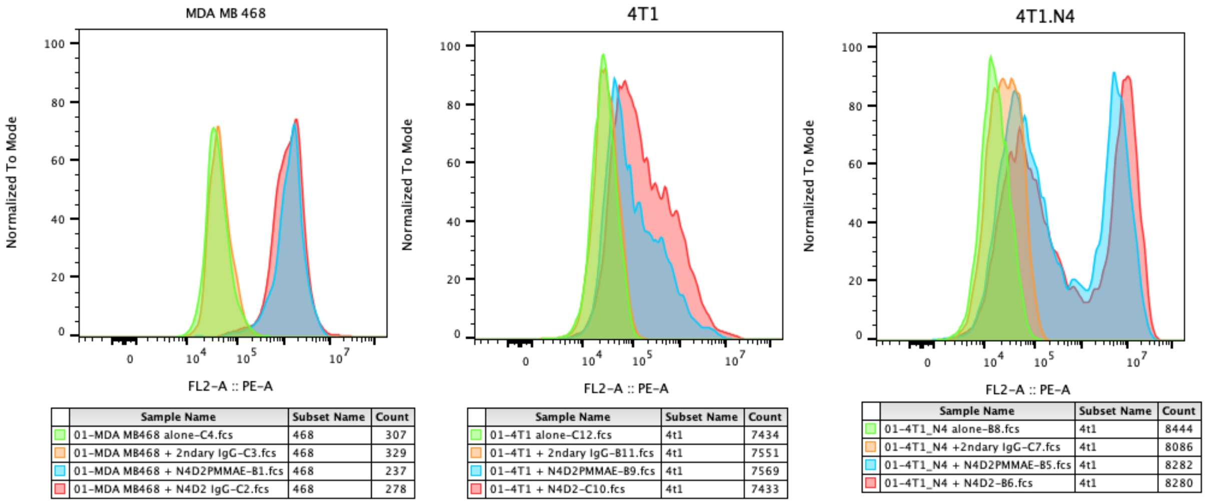


**Supplementary Figure 6**: FACS analysis of transfected 4t1 cells with human nectin-4 receptor. Different concentrations of N4MU01 were incubated with A) Nectin-4 expressing MDA-MB-468 (used as positive control), B) 4T1 cells and C) nectin-4 transfected 4T1 cells. Specific binding was observed in nectin-4 expressing MDA-MB-468 cells. Non-specific binding was observed in 4T1 cells.  In transfected 4T1 cells, mixed population of nectin-4 transfected and non-transfected 4T1 cells was observed.

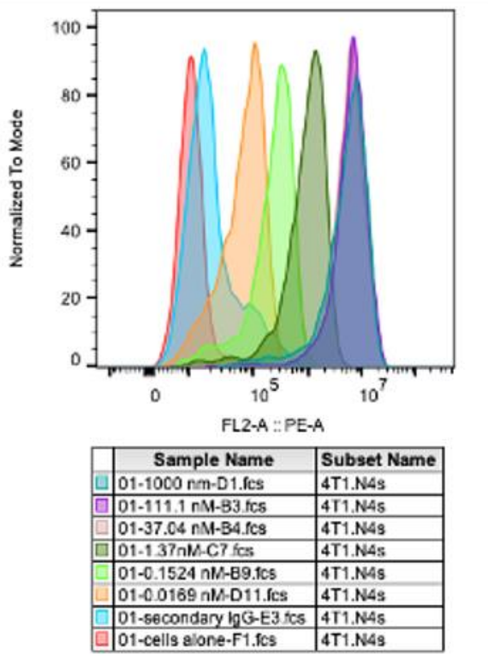


**Supplementary Figure 7**: FACS analysis of sorted nectin-4 transfected 4T1 population.

Different concentrations of N4MU01 were incubated with sorted nectin-4 transfected 4T1 cells. Dose-dependent binding was observed of the sorted 4T1 cells transfected with nectin-4.

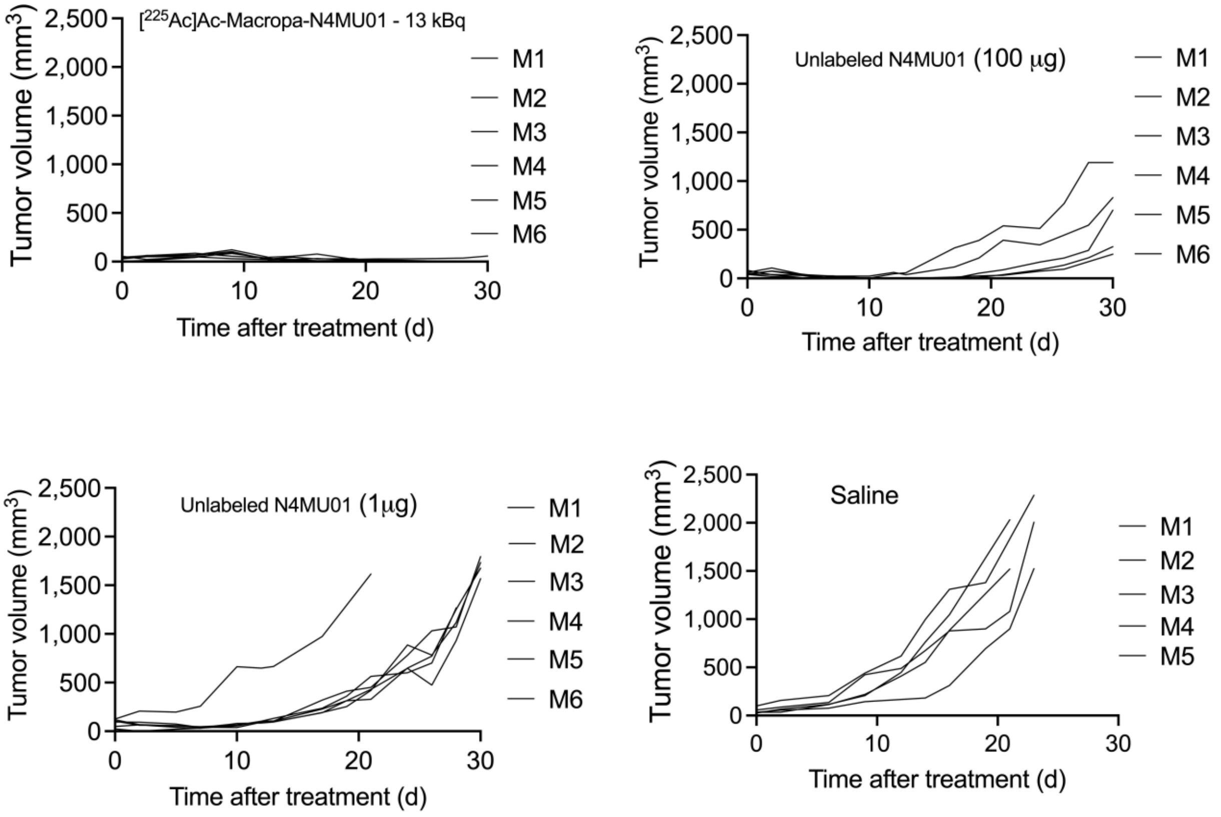


**Supplementary Figure 8:** Efficacy of [$^{225}$Ac]Ac-macropa-N4MU01 radioimmunoconjugate in 4T1.$_{\text{nectin-4}}$ syngeneic model. (A) Tumor growth curve of 13 kBq [$^{225}$Ac]Ac-macropa-N4MU01, therapeutic dose of unlabeled N4MU01 (100 μg), saline control and unlabeled N4MU01 (1.3 μg) treated mice bearing 4T1.$_{\text{nectin-4}}$ represented as tumor volumes versus days post-treatment.

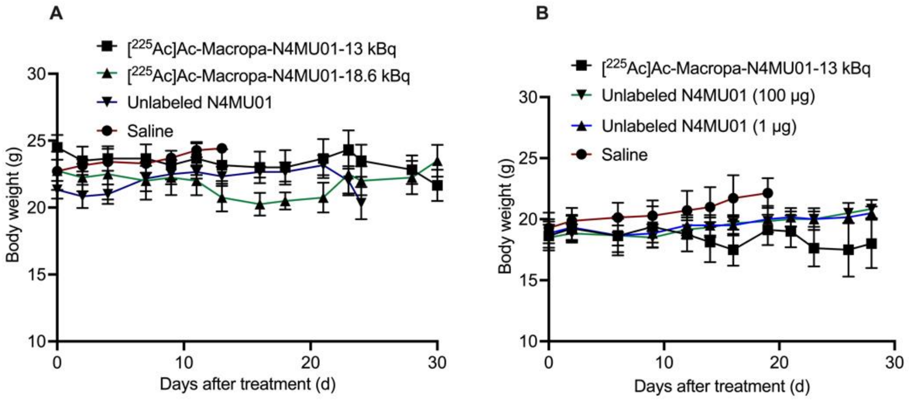


**Supplementary Figure 9:** Body weights of mice (Mean ± SEM). **A**) MDA-MB-468 (injected with Matrigel) therapy. **B**) 4T1.$_{nectin-4}$ syngeneic model therapy.
